# Single-cell, label-free morphology profiling of iPSC-derived microglia reveals dynamic state transitions

**DOI:** 10.64898/2026.04.28.721424

**Authors:** Tingting Chen, Xiaopeng Li, Amalia M Dolga

## Abstract

In response to environmental and inflammatory cues, microglia adopt diverse morphologies reflecting their functional state. However, characterizing these morphological alterations in human iPSC-derived microglia (iMGLs) remains limited by reliance on fluorescent labeling, endpoint imaging, and coarse categorical classifications (e.g., ramified vs amoeboid states). Here, we developed a label-free pipeline combining live-cell imaging, Cellpose-SAM segmentation, and CellProfiler-based feature extraction to track iMGL morphology at single-cell resolution over time. Applying this framework to a 24-hour time course of LPS and IFNγ co-stimulation revealed rapid and transient morphological remodeling, with responses peaking within 2–4 hours and partially attenuating thereafter. At single-cell resolution, four morphological states captured the major axes of variation, with co-stimulation driving a pronounced redistribution toward a single “Spread state”. Extending the analysis to individual inflammatory stimuli (LPS, IFNγ, IL-1β, IL-6, and TNFα) revealed graded but overlapping shifts in state composition, with IFNγ inducing the strongest response, whereas IL-1β showed minimal effects. Importantly, state-level changes alone were insufficient to distinguish stimulus-specific responses. Instead, single-cell density mapping revealed distinct occupancy patterns within a shared morphological landscape. In addition, texture complexity provided an independent feature layer that further separated among treatments, indicating that stimulus identity is represented by continuous, high-dimensional morphological features beyond discrete state assignments. Linking morphology to function, IFNγ produced the largest morphological redistribution and was also the only stimulus to significantly elevate intracellular ROS at 24 h, indicating that morphological remodeling and redox activation can be coordinately engaged by specific inflammatory signals. Together, these findings support a model in which microglia respond to inflammatory stimuli by graded occupancy of a continuous morphological landscape, with each stimulus producing a stimulus-specific occupancy pattern that extends beyond discrete state assignments.

## Introduction

Microglia, the brain’s main resident immune cells, originate from the embryonic yolk sac and colonize the central nervous system (CNS) during early embryonic development [1]. After migrating into the brain, microglia contribute to its development by pruning synapses, regulating neurogenesis and gliogenesis, supporting neural circuit formation and myelination, and maintaining vascular formation and integrity [2-4]. Under pathological conditions, microglia can exert both protective and detrimental effects[5]. On one hand, microglia can adopt protective roles by clearing misfolded proteins (such as amyloid-β aggregates in Alzheimer’s disease or α-synuclein in Parkinson’s disease) and cell debris. They can also release anti-inflammatory cytokines (e.g., IL-10 and TGF-β) and neurotrophic factors (e.g., BDNF and IGF-1) to support neuroprotection and regeneration. On the other hand, chronic or excessive exposure to pathological factors can drive microglia toward a detrimental inflammatory phenotype, leading to the release of cytokines such as IL-1β, IL-6, and TNF-α, which in turn trigger astrocyte activation and promote neuronal injury[6].

These functional transitions are accompanied by profound morphological remodeling, with microglia shifting from a ramified, surveying state to an amoeboid, phagocytic form as they respond to injury or disease[7]. Even under homeostatic conditions, microglia display heterogeneous morphologies, reflecting subtle regional and functional differences[8]. However, accumulating evidence indicates that microglial morphology does not conform to a simple binary classification, but instead spans a continuum of intermediate states, reflecting diverse functional programs and context-dependent responses[9, 10]. Accurate characterization of these morphological states is therefore essential for understanding microglial functional dynamics, particularly in emerging *in vitro* systems, such as induced pluripotent stem cell (iPSC)–derived microglia. These human-based models enable controlled investigation of microglial development, heterogeneity, and dynamic morphological remodeling, where cellular function is often inferred directly from morphological phenotypes[11-13].

Despite its importance, quantitative analysis of microglial morphology remains technically challenging. A defining morphological feature of microglia is their pronounced heterogeneity, even within the same physiological or pathological state[9, 14]. The complex and highly branched structure of microglia complicates accurate segmentation, especially in dense cultures with overlapping processes. Therefore, precise and unbiased quantification of microglial morphology requires accurate segmentation and cell-by-cell analysis to resolve complex cellular geometries without reliance on extensive manual intervention. To date, a range of tools have been developed to quantify microglial features such as size, ramification, soma shape, and branching architecture[15-17]. However, these pipelines typically depend on immunohistochemical labelling (e.g., Iba1, TMEM119, P2RY12, CD11b) and semi-automated tracing of high-resolution fluorescence images, which are labor-intensive and difficult to scale. Moreover, fixation and staining procedures can perturb cellular morphology, and endpoint imaging does not allow for longitudinal analysis of dynamic morphological changes[18, 19]. While genetic reporters and fluorescent probes enable live-cell imaging, they may introduce confounding effects such as phototoxicity or perturbations in their cellular processes [20-23].

Recent advances in deep learning–based image analysis have enabled robust, label-agnostic cell segmentation across diverse imaging modalities. In particular, Cellpose-SAM integrates the generalist segmentation framework with the Segment Anything Model (SAM), enabling robust and label-agnostic segmentation of cells with complex morphologies across diverse imaging conditions[24]. When coupled with scalable feature extraction platforms such as CellProfiler[25], these approaches enable high-throughput, unbiased quantification of single-cell morphological features, including cell shape, structural complexity, and spatial organization.

However, it remains unclear whether microglial inflammatory responses can be systematically encoded and resolved through quantitative morphological profiling at single-cell resolution, particularly in a dynamic, label-free setting. Furthermore, whether distinct inflammatory stimuli could induce unique morphological trajectories within a shared feature space has not yet been explored in iPSC-derived microglia.

Here, we established a high-throughput, label-free framework for mapping microglial morphological states in live iPSC-derived microglia. By integrating deep learning–based segmentation (Cellpose-SAM) with quantitative feature extraction (CellProfiler), we enable scalable and time-resolved analysis of microglial morphology without labeling. Using this approach, we demonstrated that microglial responses are continuous and stimulus-dependent and can lead to specific trajectories within a shared morphological landscape, rather than discrete binary states. This pipeline provides a quantitative and unbiased strategy to resolve microglial heterogeneity and to decode functional transition states from the analysis of the microglial morphology alone.

## Materials and methods

### Maintenance of human iPSCs and iMGL differentiation

Human Gibco iPSC line was kindly gifted by Prof. Bart Eggen from the Faculty of Medical Sciences, University Medical Center Groningen. iPSCs were maintained on Matrigel (Corning, 354277)-coated 6-well plates in E8 Flex medium (Gibco, A2858501) supplemented with E8 Flex supplement and 100 U/mL penicillin/streptomycin (Gibco, 15070063). Cells were passaged every 3–4 days using ReLeSR (STEMCELL Technologies, 100-0483) and routinely tested for mycoplasma contamination. Differentiation of iPSCs into iMGLs was performed following a previously described embryoid body–based hematopoietic differentiation protocol[26, 27], with minor modifications. Briefly, iPSCs were aggregated into embryoid bodies and sequentially differentiated through hematopoietic and myeloid progenitor intermediates using staged cytokine cocktails (BMP-4, PeproTech, 125-05; VEGF, PeproTech, 100-20; SCF, PeproTech, 300-07 for EB formation; SCF, M-CSF, PeproTech, 300-25; IL-3, PeproTech, 200-03; FLT3 ligand, PeproTech, 300-19; TPO, PeproTech, 300-18 for hematopoietic specification; FLT3 ligand, M-CSF, and GM-CSF, PeproTech, 300-03 for myeloid progenitor expansion). Microglial progenitors were collected on day 25, filtered through a 40 µm cell strainer, and matured in maturation medium consisting of Advanced DMEM/F12 (Gibco, 12634-010) supplemented with 5 µg/mL N-acetylcysteine (Sigma-Aldrich, A0737), 400 µM 1-thioglycerol (Sigma-Aldrich, M6145), 1 µg/mL heparan sulfate (Sigma-Aldrich, H7640), 1% GlutaMAX (Gibco, 35050061), 1% NEAA (Gibco, 11140-050), 1% penicillin/streptomycin, 2% B27 without vitamin A (Gibco, 17504-044), and 0.5% N2 (Gibco, 17502-048). The maturation medium was further supplemented with 100 ng/mL IL-34 (PeproTech, 200-34), 25 ng/mL M-CSF (PeproTech, 300-25), 25 ng/mL CX3CL1 (PeproTech, 300-31), and 25 ng/mL TGF-β1 (PeproTech, 100-21C). iMGLs were matured for 2 days prior to inflammatory stimulation and are referred to throughout this study as iPSC-derived microglia-like cells (iMGLs).

### Inflammatory stimulation

iMGLs were stimulated with five individual inflammatory stimuli: LPS (50 ng/mL), IFNγ (100 ng/mL), IL-1β (10 ng/mL), IL-6 (100 ng/mL), or TNFα (50 ng/mL)—or with combined LPS (50 ng/mL) and IFNγ (100 ng/mL). LPS (from *Escherichia coli*) was purchased from Sigma-Aldrich (L4391). Recombinant human IL-1β (Peprotech, 200-01B), IL-6 (Peprotech, 200-06), TNFα (Peprotech, 300-01A), and IFNγ (ImmunoTools, 11343536) were obtained through Thermo Fisher Scientific. All cytokines were reconstituted according to the manufacturer’s instructions and diluted to working concentrations in maturation medium immediately before use. Vehicle controls received an equivalent volume of cytokine reconstitution buffer (0.1% BSA in sterile water) to match carrier protein content across all conditions. Three independent differentiation runs were conducted as biological replicates (Exp1, Exp2, Exp3). All analyses presented in this study use the IL-34-supplemented condition; the IL-34– condition was acquired in parallel as part of a separate experimental arm and is reported elsewhere.

### Live-cell imaging

Immediately after stimulation, plates were transferred to an IncuCyte S3 live-cell imaging system (Sartorius) and maintained at 37°C with 5% CO_2_. Phase-contrast images were acquired every 2 h for 24 h at 10× magnification, with 5 fields of view per well. Two fields (fields 1 and 3) were selected for downstream analysis to sample well coverage while minimizing spatial autocorrelation between images of the same well.

### Image preprocessing

Prior to segmentation, raw phase-contrast images were preprocessed in Python using PIL (Pillow v 11.3.0). Multi-page TIFFs were reduced to their first frame and converted to 8-bit grayscale. Each image was then sharpened with an unsharp mask (radius = 10 pixels, percent = 100, threshold = 0) followed by a Gaussian blur (radius = 0.4 pixels) to enhance cell boundaries while suppressing high-frequency noise. The full preprocessing script is provided in the accompanying GitHub repository.

### Image segmentation and feature extraction

Preprocessed images were segmented at single-cell resolution using Cellpose-SAM v4.0.6[24] running on Python 3.12.3 with PyTorch 2.5.1 (CUDA 12.1). Segmentation was performed using the pretrained cpsam model on GPU without fine-tuning. Images were processed as single-channel grayscale input (channels = [0, 0]). A fixed object diameter of 25 pixels was used across all images. Segmentation parameters were empirically adjusted to optimize performance for this dataset (flow_threshold = 1.1, cellprob_threshold = −3), and image normalization was applied using percentile scaling (1st–99th percentile). Inference was run with 600 iterations and a batch size of 1. Quantitative morphological features were extracted from Cellpose-SAM segmentation masks using CellProfiler *v4*.*2*.*8*[25]. The pipeline computed per-cell shape descriptors (MeasureObjectSizeShape), spatial neighbor relationships (MeasureObjectNeighbors), and Haralick texture features on the raw phase-contrast images (MeasureTexture) and exported per-cell measurements as CSV files via ExportToSpreadsheet. Of the features extracted, eight (Area, Perimeter, Solidity, Extent, Eccentricity, FormFactor, NumberOfNeighbors_Adjacent, PercentTouching_Adjacent) were retained for downstream analysis. Texture features were used to derive a continuous texture complexity score (defined below); they were not included in the feature set used for state assignment or for the UMAP embedding.

### Non-cell object filtering

Within each experiment, objects were retained if area ≥ 50 pixels^2^, log(area) lay within ±5 median absolute deviations of the per-experiment median, and Solidity ≥ 0.70. This typically retained 90–95% of segmented objects per experiment.

### Image-level aggregation and Vehicle-based per-experiment centering

Per-cell features were aggregated to image-level medians; images with fewer than 20 cells were excluded. Area, Perimeter, NumberOfNeighbors_Adjacent, and PercentTouching_Adjacent were log1p-transformed. Vehicle values were first averaged within each time point, then combined by unweighted mean across time points to yield one offset per experiment. Each feature was centered by subtracting its experiment’s offset. Global PCA on the centered matrix (prcomp, center = TRUE, scale. = TRUE) provided the coordinate system for all image-level analyses.

### Single-cell pre-processing

Single-cell analyses in the main figures use two time points: 4 h (early remodelling phase) and 24 h (sustained response phase), chosen independently of this dataset on the basis of commonly used time windows in the microglial cytokine-response literature[27, 28]. For single-cell state analyses, cells at the pre-specified time points were log-transformed and centered with the same experiment-level Vehicle offset as the image-level data. Cells were then subsampled to 1,667 per (Stimulus × Experiment) by stratified random sampling (seed = 123); when a group contained fewer cells, the sample size was reduced to the minimum group size.

### UMAP embedding

The centered, z-scored single-cell matrix was reduced by PCA, retaining the smallest number of components explaining ≥95% of cumulative variance. UMAP was computed on these PCs (uwot::umap; n_neighbors = 50, min_dist = 0.5, metric = “euclidean”, seed = 123). No post-hoc coordinate filtering was applied. UMAP was used exclusively for visualization; all statistical tests used the complete (non-subsampled) single-cell data.

### Morphological state anchors

Canonical morphological states were defined once, on a fixed training set, and then applied to all downstream figures by nearest-anchor assignment. The training set comprised all cells from the Vehicle and the co-treatment of LPS and IFNγ conditions, pooled across all 13 time points and three experiments. After log-transformation and experiment-level centering, cells were projected onto three z-scored axes — z_size (log Area), z_round (FormFactor), and z_elong (Eccentricity) — using scaling parameters (mean, SD) computed on this training set. K-means (k = 4, nstart = 25, iter.max = 50, seed = 123) was fit in this three-axis space. Clusters were labelled deterministically: the cluster with the highest z_elong centroid was named Elongated; of the remaining three, the highest z_size was Spread; of the remaining two, the highest z_round was Small_Round; the last was Ramified. The four anchor centroids and the scaling parameters were saved (R/state_anchors.rds) and reused in every downstream figure. As a sensitivity analysis for the choice of k, mean silhouette widths were computed for k = 2–8 on a 5,000-cell random subsample of the training set. The k = 4 solution was selected to balance geometric separation with interpretability and morphological granularity, because lower-k solutions merged multiple final anchor states that were separable in the final anchor space (Fig. S3). In all downstream analyses, each cell was assigned to the state whose anchor was closest in squared Euclidean distance in the frozen z-axis space, after applying the saved scaling parameters. No re-clustering or cross-figure cluster matching was performed.

### Texture complexity

Texture complexity was defined as PC1 of 13 direction-averaged Haralick features extracted by CellProfiler (MeasureTexture, scale = 10 pixels, 256 gray levels). For each Haralick descriptor (AngularSecondMoment, Contrast, Correlation, DifferenceEntropy, DifferenceVariance, Entropy, InfoMeas1, InfoMeas2, InverseDifferenceMoment, SumAverage, SumEntropy, SumVariance, Variance), the four directional measurements (0°, 45°, 90°, 135°) were averaged to yield a rotationally invariant value per cell. The 13 features were z-scored and subjected to PCA. PC1 explained approximately 50.4% of the total variance and was oriented so that higher values correspond to higher SumEntropy. For visualization, values were clipped at the 1st and 99th percentiles. Texture complexity was not used in the UMAP embedding or in state assignment.

### Statistical analysis

All hypothesis tests were performed at the experiment level (n = 3 independent differentiation runs) to avoid pseudoreplication: single-cell and image-level quantities were aggregated to per-(Stimulus × Experiment) summaries before testing. Tests were two-sided. Where multiple comparisons were performed within a panel, p-values were adjusted by the Benjamini–Hochberg procedure with the correction scope reported in the corresponding figure legend. Linear mixed-effects models (Fig. 4E,F; Fig. 6B) were fit with lme4 and lmerTest, with stimulus-vs-Vehicle contrasts obtained by Dunnett’s test with multivariate-t adjustment (emmeans). The specific test, sample size, and FDR scope used for each comparison are stated in the corresponding figure legend.

### Code availability

Downstream analyses were performed in R v4.4.2 using tidyverse, uwot, patchwork, ggpubr, ggforce, ggrepel, cluster, MASS, scales, lme4, lmerTest, and emmeans. Segmentation used Cellpose-SAM v4.0.6 running on Python 3.12.3 with PyTorch 2.5.1 and CUDA 12.1; image preprocessing used Pillow v11.3.0; and feature extraction used CellProfiler v4.2.8. Random seeds were fixed at 123 for all stochastic steps. Complete analysis code, image preprocessing and segmentation scripts, CellProfiler pipelines, metadata files, and the frozen anchor file (R/state_anchors.rds) are available at https://github.com/TingtingChen311/microglia-morphology-profiling and archived on Zenodo at https://doi.org/10.5281/zenodo.19826203.

## Results

### 1. A label-free pipeline for single-cell morphological profiling of human iPSC-derived microglia

To facilitate characterization of microglial morphology in a scalable, unbiased manner in human iPSC-derived microglia (iMGLs), we established an integrated, label-free pipeline. This strategy combined live-cell time-lapse imaging, deep learning–based segmentation, and quantitative single-cell feature extraction (Fig. 1A). iMGLs were seeded in 96-well plates and imaged under phase contrast every 2 h for 24 h using an Incucyte live-cell imaging system. The morphology of single cells was then segmented using Cellpose-SAM, which reliably resolved densely packed iMGLs with overlapping processes across imaging fields (Fig. 1B). From each segmented cell, we extracted a panel of shape and spatial features using CellProfiler. Using these combined programs, we could generate high-dimensional single-cell morphological profiles suitable for downstream trajectory mapping and microglial state characterisation.

**Figure 1.**
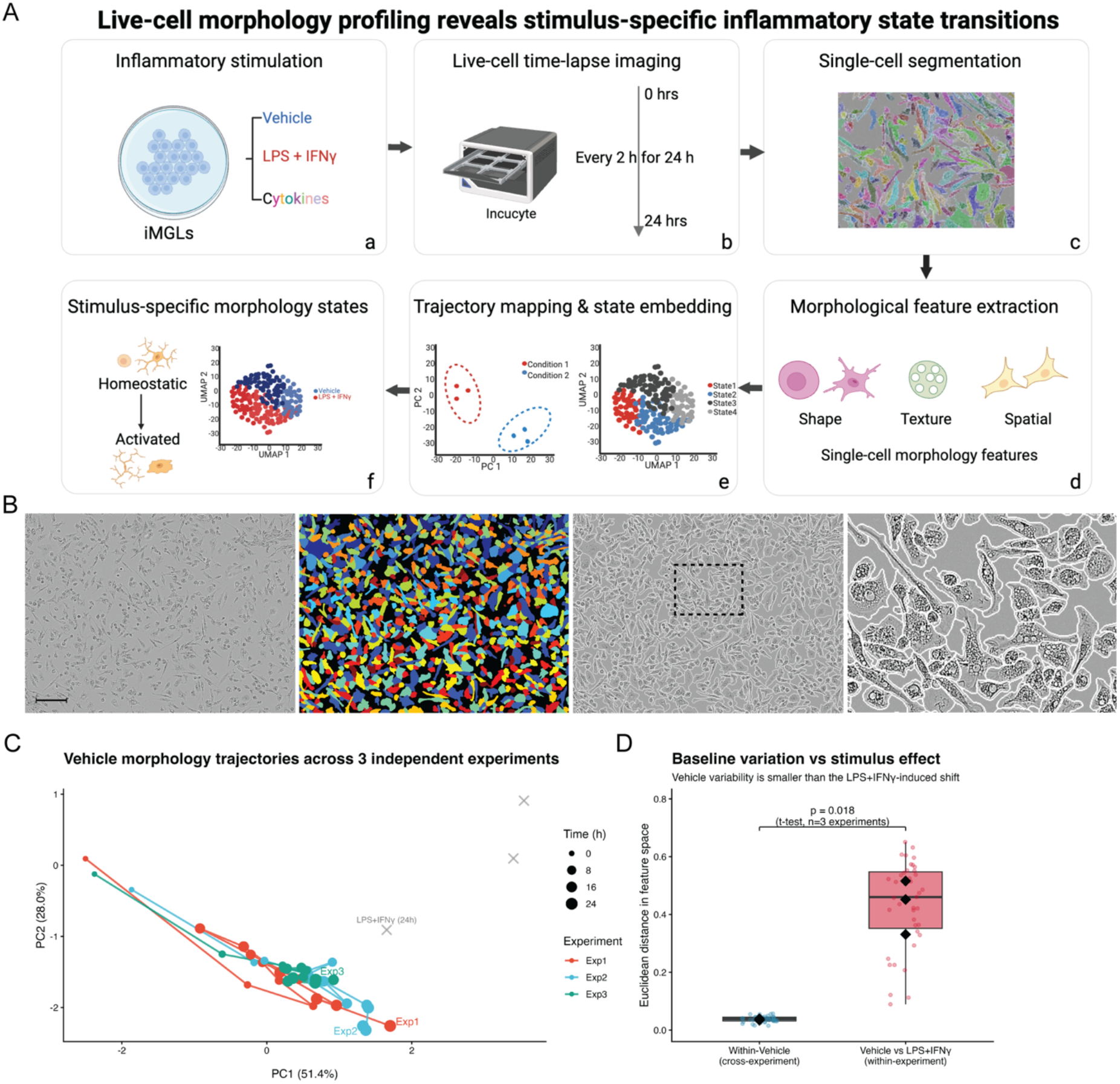
Workflow and validation of the label-free morphological profiling pipeline. (A) Workflow overview: iMGLs were stimulated with vehicle, LPS and IFNγ (depicted as LPS+IFNγ in the graph), or individual inflammatory stimuli, imaged every 2 h for 24 h using an Incucyte S3 system. Images were segmented using Cellpose-SAM, followed by single-cell feature extraction with CellProfiler for downstream PCA and UMAP analyses. (B) Representative segmentation example showing a phase-contrast image (scale bar, 200 µm), corresponding to a Cellpose-SAM mask, a zoomed inset, and a segmentation overlay. (C) PCA trajectories of vehicle-treated iMGLs across three independent experiments. Point size indicates time progression, and gray crosses represent LPS and IFNγ-treated cells at 24 h exposure. (D) Euclidean distances in morphological feature space. Left: pairwise distances between vehicle samples from different experiments (cross-experiment baseline). Right: distances between vehicle and LPS+IFNγ samples within the same experiment, *p* = 0.018, two-sided Welch’s *t*-test, n = 3 per group (experiment-level means).

To assess whether this pipeline can robustly distinguish biological effects from technical variation, we profiled vehicle-treated iMGLs across three independent experiments and compared cross-experiment variability within the vehicle condition to treatment-induced differences within experiments. Overall, in the global PCA space, vehicle-treated iMGLs from three independent experiments followed highly consistent trajectories and clustered tightly together (Fig. 1C). iMGLs treated with combined LPS and IFNγ for 24 h occupied a clearly distinct region of the PCA space, indicating a strong treatment-induced shift. Consistently, cross-experiment variability within the vehicle condition was significantly smaller than treatment-induced differences (*p* = 0.018, Fig. 1D), indicating that biological signal exceeded technical variation. These findings validated the reproducibility of the proposed framework for resolving microglial morphology at single-cell resolution over time.

### 2. Time-lapse profiling captures the dynamic morphological responses of iMGLs to LPS and IFNγ

To validate that our pipeline could capture alterations in microglial morphological dynamics, we examined the response of iMGLs to canonical proinflammatory stimuli, LPS and IFNγ, following a 24 h exposure. Phase-contrast imaging showed a clear divergence between vehicle- and co-stimulated cells, as early as 2 h and even more pronounced at 4 h following treatment. Co-stimulated cells with LPS and IFNγ showed a more compact, less ramified morphological profile compared to vehicle controls (Fig. 2A). PCA across all time points quantitatively confirmed this divergence. At 0 h, vehicle and co-stimulated cells largely overlapped, but progressively separated over time, with maximal separation observed at 4 h (Fig. 2B).

**Figure 2.**
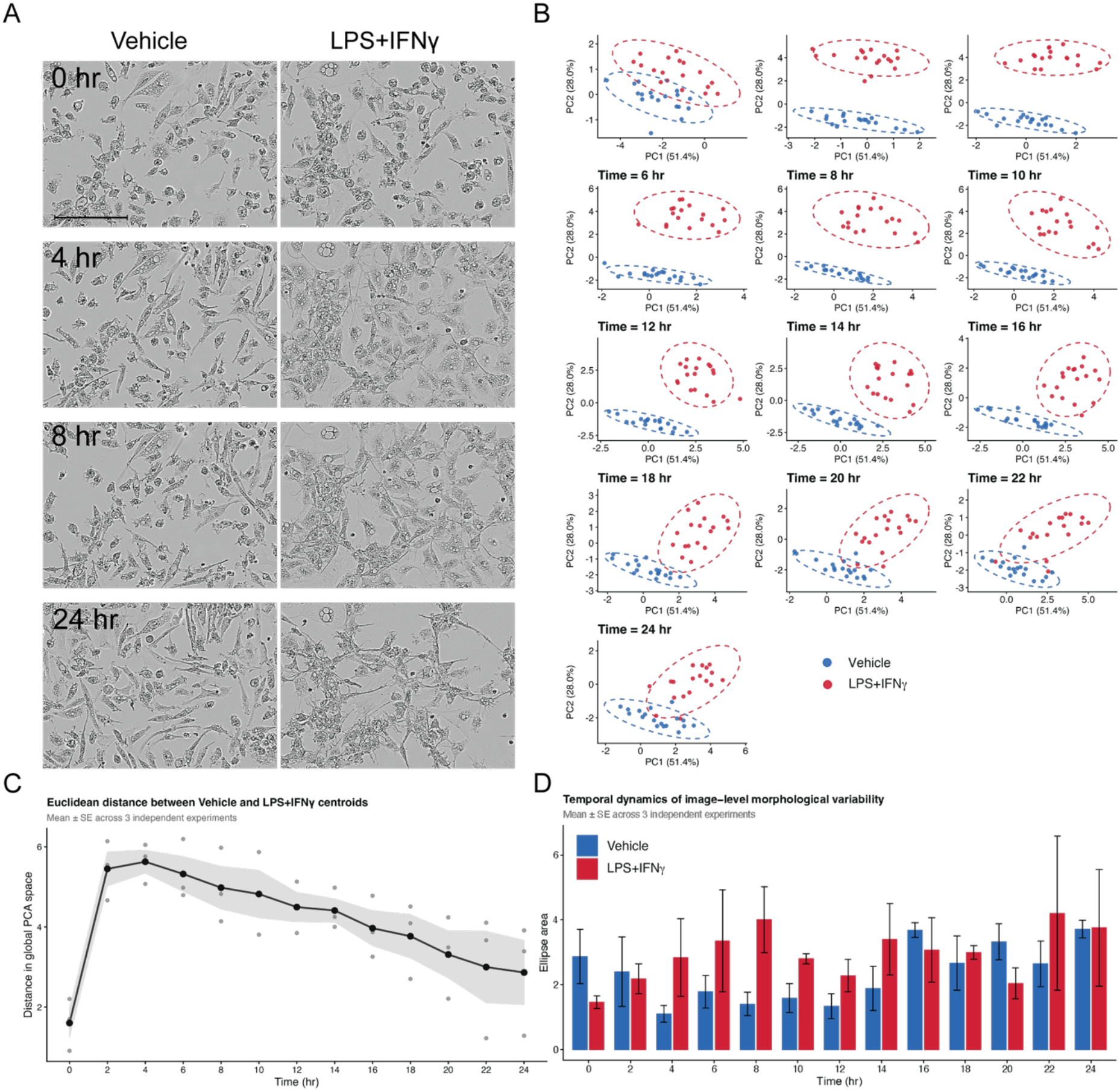
Time-resolved morphological response of iMGLs to combined treatment of LPS and IFNγ over 24 h. (A) Representative phase-contrast images of vehicle and co-stimulated iMGLs at 0, 4, 8, and 24 h. Scale bar = 200 µm. (B) PCA of image-level morphological profiles at each imaged time point. Dashed ellipses show 68% confidence intervals per treatment. (C) Euclidean distance between vehicle and co-stimulation centroids in global PCA space over time. Mean ± SE across three independent experiments; gray dots indicate per-experiment values. (D) Morphological heterogeneity quantified as the area of the 95% confidence ellipse in the PCA space. Bars show mean ± SE across three independent experiments.

Notably, the Euclidean distance between vehicle and co-stimulation centroids followed a transient trajectory in the global PCA space, increasing sharply between 2 and 4 h, followed by a gradual decline toward the baseline over the subsequent 20 h (Fig. 2C). Meanwhile, the heterogeneity in the iMGLs morphology, quantified as the area of the 95% confidence ellipse in the PCA space, was generally higher in co-stimulation group compared to vehicle, particularly between 4 and 14 h (Fig. 2D). These results show that live-cell morphological profiling captures both the magnitude and the temporal dynamics of the iMGL response to LPS and IFNγ. Overall, this analysis could reveal transient phenotypic states that would otherwise be obscured in endpoint analyses.

### 3. Single-cell embedding resolves four distinct morphological states

Initial analysis revealed robust population-level shifts induced by LPS and IFNγ co-stimulation. However, it remained unclear whether these changes reflected uniform remodeling across all cells or selective enrichment of distinct morphological subpopulations. To address this, we projected segmented single cells at two pre-specified time points, 4 h (early response) and 24 h (sustained response), into a shared UMAP embedding built from shape and spatial features. Nearest-anchor assignment using frozen reference profiles identified four morphological states, which we termed “Elongated”, “Spread”, “Ramified”, and “Small_Round”, based on their characteristic feature profiles and visual inspection of representative cells (Fig. 3A, B). Elongated cells featured high eccentricity and low form factor; Spread cells were defined by large area and high solidity; Ramified cells showed intermediate size with irregular morphology; and Small_Round cells were compact and round (Fig. 3B).

**Figure 3.**
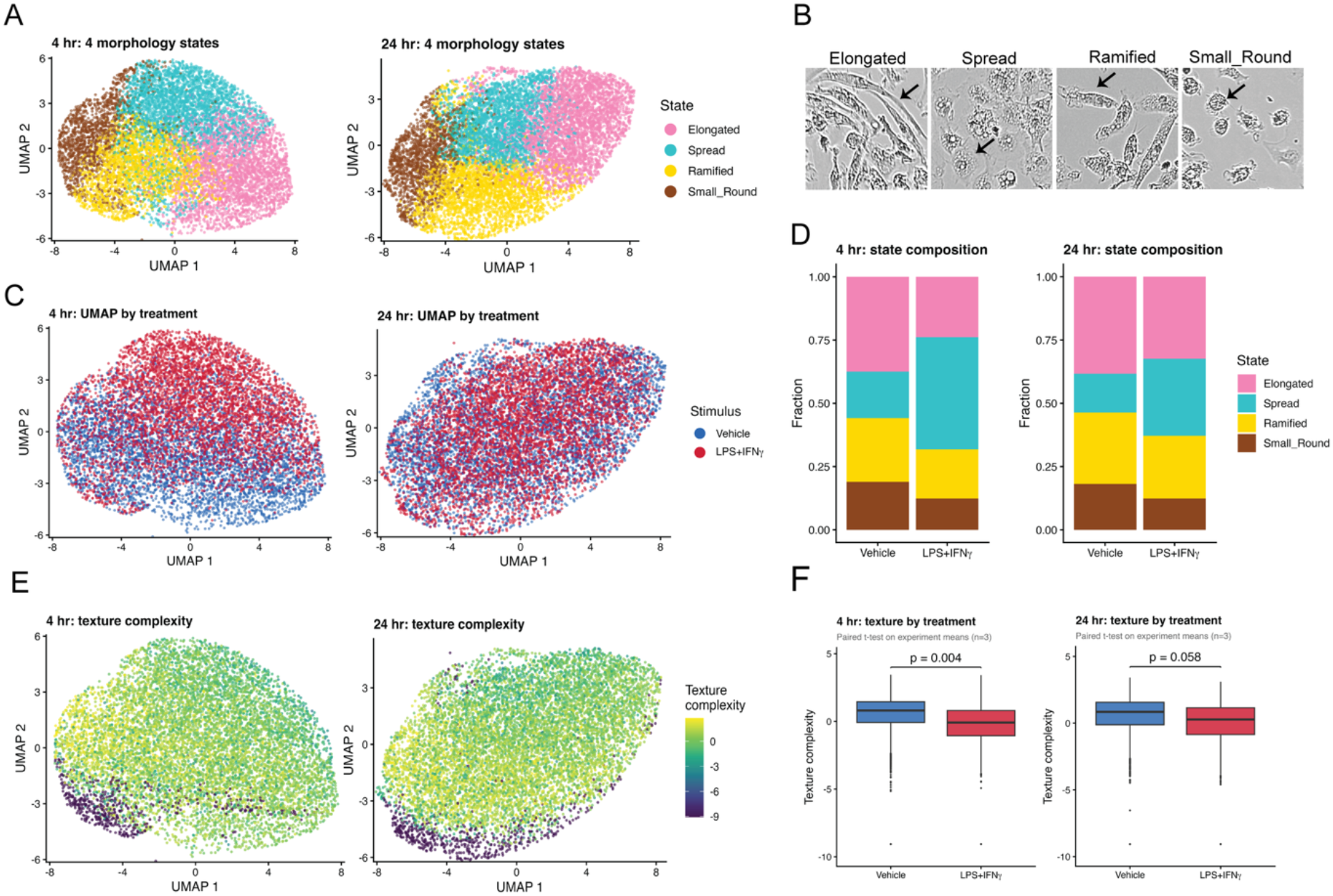
UMAP-based identification and characterization of four morphological states. (A) UMAP embedding of single iMGLs at 4 h (left) and 24 h (right), pooled across vehicle and co-stimulation conditions, colored by morphological state assigned via nearest-anchor classification in a frozen three-axis state space. (B) Representative phase-contrast images illustrating each state; black arrows indicate cells representative of the labeled state. (C) Same UMAP as in (A), colored by treatment (blue, vehicle; red, LPS+IFNγ). (D) State composition per treatment at 4 h and 24 h. Statistical testing was performed using paired t-test on per-experiment state fractions (n = 3 independent experiments); BH-FDR within each time point across the 4 morphological states; see Table S1. (E) Same UMAP as in (A), colored by texture complexity (PC1 of 13 direction-averaged Haralick features; higher values = higher complexity). (F) Texture complexity by treatment at 4 h and 24 h. Per-experiment image-level texture-complexity medians were compared between vehicle and co-stimulation of LPS and IFNγ by paired *t-*test (n = 3 independent experiments); no multiple-testing correction was applied as only two comparisons were tested. Box plots show median, IQR, and 1.5×IQR whiskers; points beyond the whiskers are individual cells.

**Figure 4.**
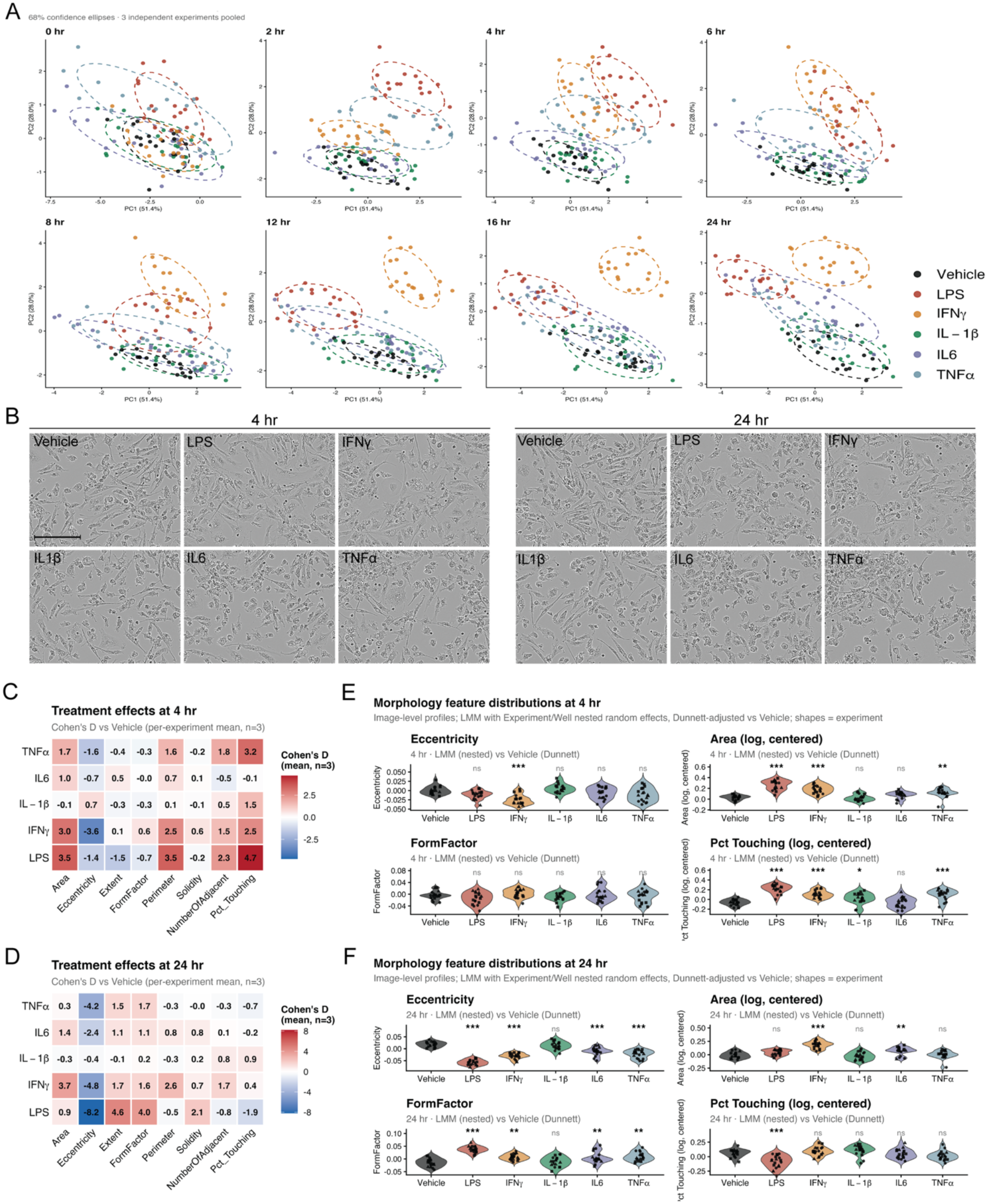
Image-level morphological effects of five inflammatory stimuli across time. (A) PCA of image-level morphological profiles across 13 time points (0–24 h) for iMGLs treated with vehicle, LPS, IFNγ, IL-1β, IL-6, or TNFα; for visual clarity, panel A shows 8 representative time points. (B) Representative phase-contrast images at 4 h (left) and 24 h (right) for each treatment. Scale bar: 200 µm. (C, D) Cohen’s D of each morphological feature versus vehicle at 4 h (C) and 24 h (D). Significance was assessed by one-sample t-test on the three per-experiment D values (H_0_: mean D = 0), with BH-FDR correction across all feature × stimulus comparisons within each time point (*FDR < 0.05, **FDR < 0.01, ***FDR < 0.001). (E, F) Image-level distributions of four representative features at 4 h (E) and 24 h (F). Points: individual images (3 wells × 2 images per condition per experiment; n = 3 experiments; shapes denote experiment). Significance: linear mixed-effects model on the centered, log-transformed image-level values (Area, Perimeter log-transformed; Eccentricity, FormFactor untransformed; see Methods), with nested random intercepts for Experiment and Well; stimulus-vs-Vehicle contrasts obtained by Dunnett’s test with multivariate-t adjustment (n = 3 wells × 2 images × 3 experiments per condition). *\*p* < 0.05, *\*\*p* < 0.01, *\*\*\*p* < 0.001; ns, not significant.

When split by treatment, vehicle-treated cells were distributed across all four states, whereas co-stimulation led to a clear shift toward the Spread state, with the strongest effects at 4 h (Fig. 3C, D). At 4 h, paired *t*-tests on per-experiment state fractions confirmed that Spread was strongly enriched (Δ = +0.26, FDR = 0.013) while Elongated, Ramified, and Small_Round were correspondingly reduced (all FDR < 0.05; Table S1, BH-FDR within time point across the 4 states). By 24 h, these state shifts had partially attenuated: the Spread state remained elevated in effect size (Δ = +0.15) but did not reach FDR significance after correction (FDR = 0.087); see Fig. S1A for the full time course.

To further characterize these states, we overlaid an independent texture-complexity feature onto the same embedding. Texture values showed a clear gradient matching the state structure: Ramified and Small_Round cells generally showed higher texture complexity, while Elongated and Spread cells showed lower complexity (Fig. 3E). Consistent with the enrichment of the low-complexity Spread state, co-stimulation reduced overall texture complexity at 4 h (*p* = 0.004), with the effect diminishing by 24 h (*p* = 0.058; Fig. 3F). Because texture features were not used to define the states, these results indicate that the four morphological states capture coherent, multi-dimensional differences in cell structure that extend beyond the shape and spatial features used to define them.

### 4. Inflammatory stimuli induce time-dependent morphological signatures in iMGLs

Having established the pipeline on a canonical stimulus, we next asked whether exposure to individual inflammatory stimuli could induce distinct and time-dependent morphological signatures. iMGLs were treated with vehicle, LPS, IFNγ, IL-1β, IL-6, or TNFα and we next performed time-lapse profiling across 24 h. Global PCA of image-level profiles revealed that all six conditions overlapped at 0 h but progressively diverged over time, with a separation becoming apparent at 2–4 h and persisting through the next 16 h (Fig. 4A). Phase-contrast images at 4 h and 24 h reflected these morphological changes (Fig. 4B).

To quantify these effects, we computed per-experiment Cohen’s D relative to vehicle (paired within experiment) and averaged across 3 independent experiments (Fig. 4C, D). Significance was assessed by one-sample *t*-test on the three per-experiment D values, with Benjamini–Hochberg FDR correction within each time point. At 4 h, LPS and IFNγ produced the strongest effects across multiple features, including increased Area and Perimeter and reduced Eccentricity (LPS: D_Area = 3.5, D_Eccentricity = −1.4; IFNγ: D_Area = 3.0, D_Eccentricity = −3.6). LPS additionally reduced extent (D = −1.5) and strongly increased cell–cell contact (Pct_Touching, D = 4.7), while IFNγ increased FormFactor (D = 0.6) and Solidity (D = 0.6). IL-1β, IL-6, and TNFα induced weaker and more limited changes at 4 h (Fig. 4C).

At 24 h, both the magnitude and pattern of responses had shifted. IFNγ showed strong effects across a broader set of features (D_Area = 3.7, D_Eccentricity = −4.8, D_Extent = 1.7, D_FormFactor = 1.6, D_Perimeter = 2.6) but only a small effect on Pct_Touching (D = 0.4). LPS primarily affected shape, with a markedly smaller effect on Area (D = 0.9 vs 3.5 at 4 h) and a reversal of its Pct_Touching effect (D = −1.9 at 24 h vs +4.7 at 4 h). TNFα showed a pattern dominated by strongly reduced Eccentricity (D = −4.2) with intermediate effects on Extent and FormFactor, while IL-6 showed milder but broadly similar directional changes. IL-1β remained the least responsive across both time points (Fig. 4D). Image-level distributions of representative features at both time points confirmed these treatment-specific effects (Fig. 4E, F). Together, these results demonstrate that each proinflammatory stimulus elicits a distinct and time-dependent morphological signature.

### 5. Single-cell morphology reveals stimulus-specific signatures within a shared morphological landscape

Image-level profiling (Fig. 4) demonstrated that each inflammatory stimulus produces a distinct average morphological response but could not resolve whether these effects reflect shifts among pre-existing cell populations or the emergence of treatment-specific states. To address this, we applied the four-state framework defined in Fig. 3 to single cells from the five-stimulus experiment, focusing on 4 h (early response) and 24 h (sustained response). Post-hoc inspection of global PCA separation (Fig. 4A) and state-fraction dynamics (Fig. S2) confirmed that these time points capture, respectively, the early peak and the sustained-response phase of the morphological remodeling.

At each time point, a matched number of cells per condition (n = 1,667 cells × 6 conditions × 3 experiments) was projected into a shared UMAP embedding. The four morphological states were clearly separated in UMAP space at both time points (Fig. 5A, C), and all six treatments occupied a largely overlapping landscape, indicating that inflammatory stimulation does not generate novel, treatment-specific morphological states.

**Figure 5.**
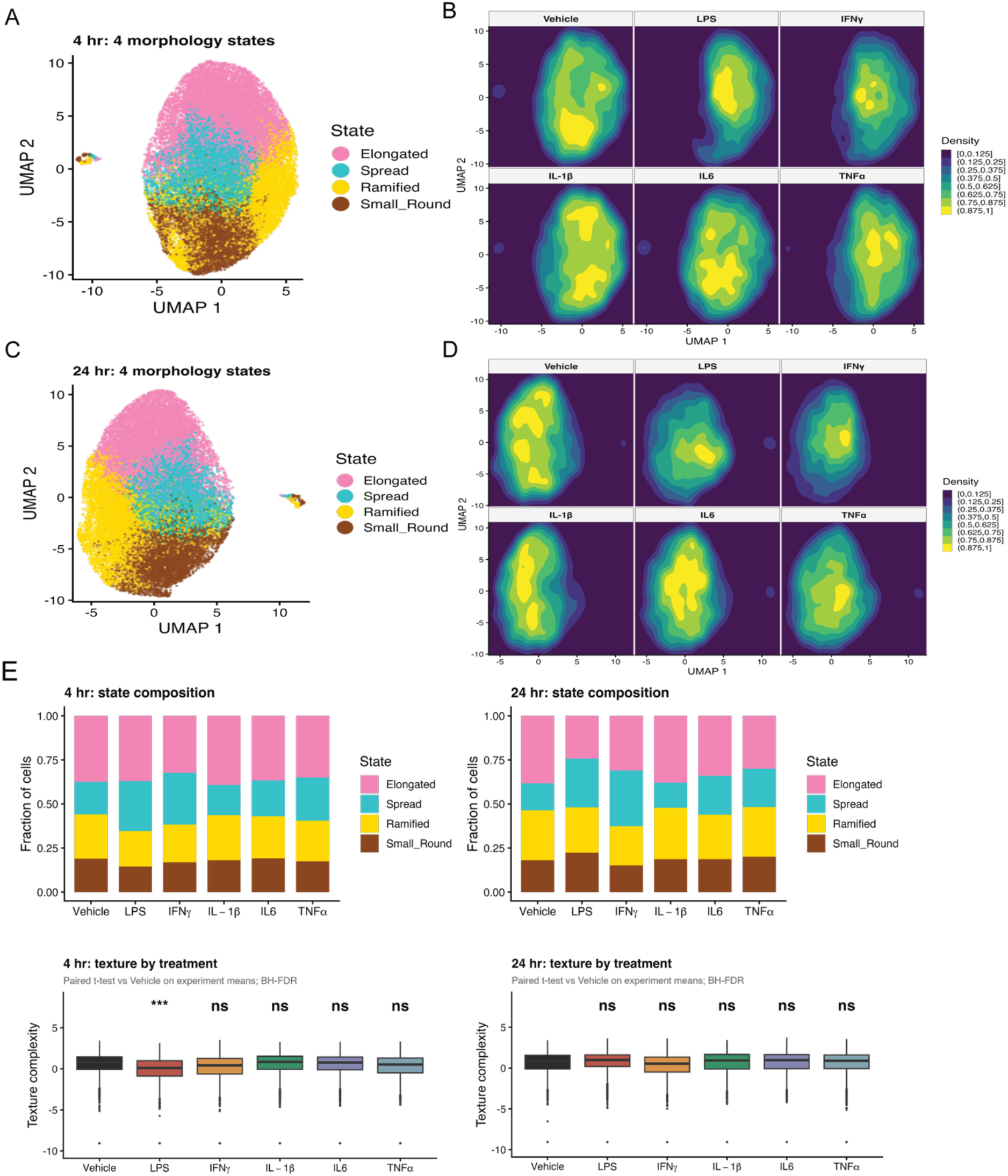
Single-cell morphological landscapes across five inflammatory stimuli at 4 h and 24 h. (A) UMAP embedding of single iMGLs at 4 h, pooled across vehicle, LPS, IFNγ, IL-1β, IL-6, and TNFα, colored by morphological state. (B) Per-treatment density maps in the same UMAP space at 4 h; density is normalized within each treatment to highlight relative occupancy patterns. (C) UMAP embedding at 24 h, colored by morphological state. (D) Per-treatment density maps at 24 h. (E) State composition (fraction of cells per state) within each treatment at 4 h and 24 h. Bars show mean ± SE across n = 3 independent experiments. Statistical testing was performed at the experiment level (paired t-test on per-experiment state fractions; BH-FDR within each time point across all 4 states × 5 stimulus-vs-vehicle comparisons (20 tests per time point)); see Table S2. (F) Texture complexity (PC1 of 13 direction-averaged Haralick features) by treatment at 4 h and 24 h. Per-experiment image-level texture-complexity medians were compared between each stimulus and vehicle by paired t-test (n = 3 independent experiments); BH-FDR correction was applied within each time point across the 5 stimulus-vs-vehicle comparisons.

**Figure 6.**
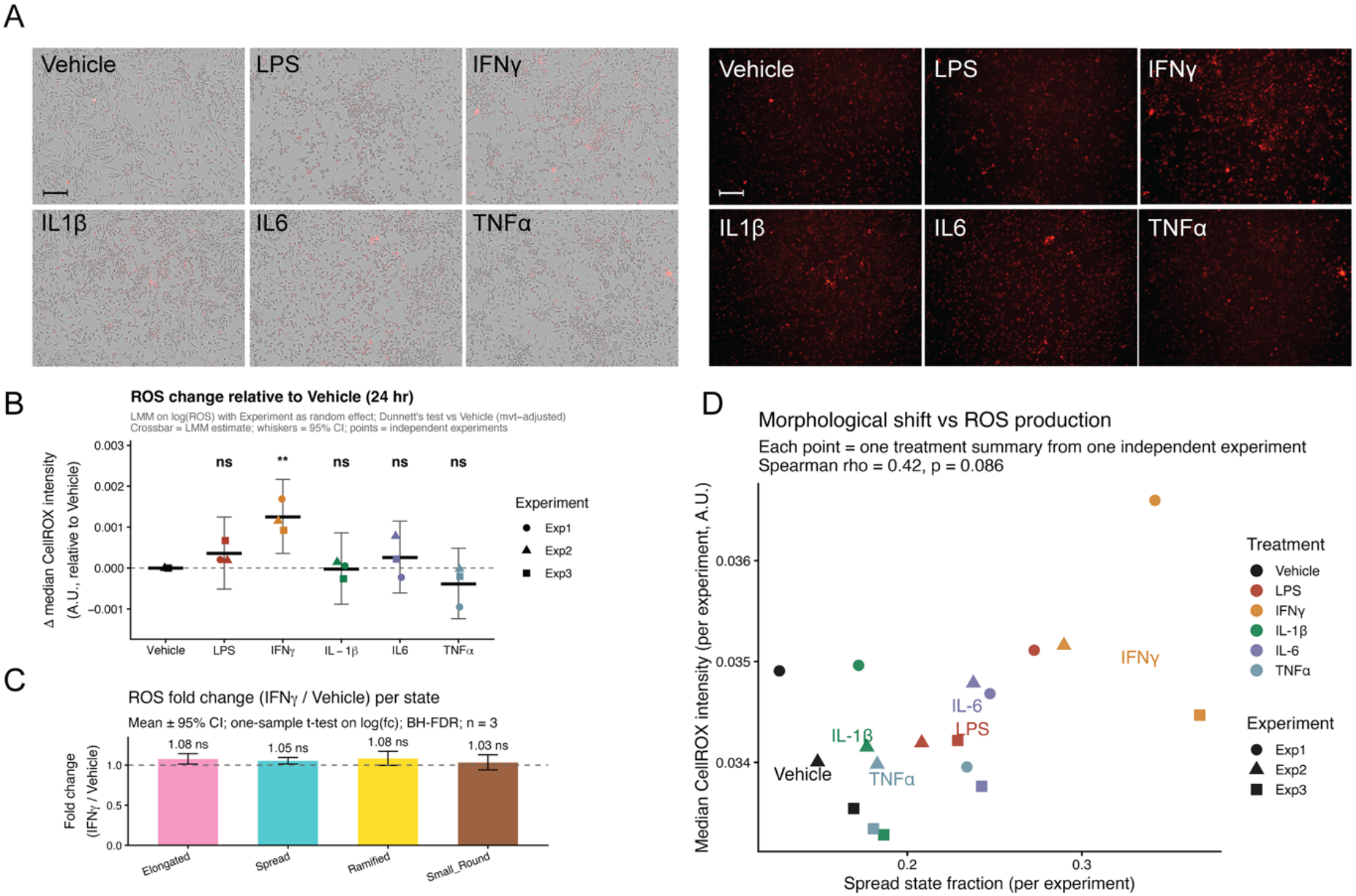
IFNγ specifically elevates intracellular ROS at 24 h and correlates with Spread-state enrichment. (A) Representative phase-contrast (left) and CellROX deep-red fluorescence (right) images of iMGLs after 24 h inflammatory stimulation. Scale bar, 200 µm. (B) Change in per-cell median CellROX intensity relative to vehicle. Points: per-experiment medians of image-level CellROX intensity (n = 3 independent experiments); crossbars and whiskers: LMM-estimated mean and 95% CI. Linear mixed-effects model on log-transformed image-level CellROX medians with Experiment as a random intercept; stimulus-vs-Vehicle contrasts obtained by Dunnett’s test with multivariate-t adjustment. *\*\*p* < 0.01; ns, not significant. (C) IFNγ/Vehicle ROS fold change per morphological state. For each experiment and each state, the per-cell CellROX median was computed for IFNγ and for Vehicle, the IFNγ/Vehicle ratio was log-transformed, and the resulting log fold-changes (n = 3 independent experiments) were tested against zero by one-sample *t*-test; BH-FDR correction was applied across the four morphological states. Bars: geometric mean across experiments; error bars: 95% CI. (D) Per-experiment Spread-state fraction vs median CellROX intensity. Each point is one (treatment × experiment) combination (6 treatments × 3 independent experiments, n = 18). Spearman’s rank correlation; *ρ* and *p*-value reported on the panel.

Per-treatment density maps revealed graded but visible stimulus-specific shifts within this shared landscape (Fig. 5B, D). At 4 h, the six conditions showed subtle offsets in the position of their density peaks. By 24 h, differences were more pronounced: IFNγ appeared visually most distinct from vehicle, with depletion of the central high-density region and a shifted hotspot; LPS also deviated clearly; IL-1β remained closest to vehicle.

At 4 h, state shifts were modest across all treatments, with no contrast reaching FDR < 0.05 after Benjamini-Hochberg correction within the time point (across 20 state × stimulus comparisons; Table S2). The closest to significance were LPS-induced reductions in the Ramified and Small_Round fractions (both Δ ≈ −0.06, raw *p* < 0.01, FDR = 0.067). At 24 h, IFNγ produced the largest effect-size shifts among the six conditions, with an increase in the Spread fraction (Δ = +0.16, raw *p* = 0.015, FDR = 0.074) and decreases in Elongated (Δ = −0.09, FDR = 0.12) and Ramified (Δ = −0.05, FDR = 0.13) states. LPS produced shifts of comparable magnitude in the same direction (Spread Δ = +0.13, FDR = 0.074; Elongated Δ = −0.16, FDR = 0.074) and was the only stimulus with a positive shift toward the Small_Round state (Δ = +0.04, raw *p* = 0.064, FDR = 0.15). TNFα produced smaller shifts of the same sign. IL-6 produced the only contrast that reached FDR significance: a modest but highly reproducible increase in the Spread fraction at 24 h (Δ = +0.07, FDR = 0.009), reflecting low between-experiment variance across the three independent replicates. IL-1β did not differ from vehicle in any state at either time point. Overall, after BH-FDR correction within each time point, only IL-6 Spread at 24 h crossed the conventional FDR < 0.05 threshold; other shifts, despite being larger in magnitude, did not pass the conservative correction with n = 3 biological replicates. Critically, all treatments retained cells in all four states, indicating that stimulus identity modulates the balance among morphological states rather than generating new ones.

To capture morphological differences beyond discrete state classification, we defined a continuous “texture complexity” score as PC1 of direction-averaged Haralick features (Fig. 5F). At 4 h, LPS produced a strong reduction in texture complexity relative to vehicle (paired *t*-test on per-experiment image medians, BH-FDR = 6.8 × 10^−4^ within time point); IFNγ showed a borderline reduction (FDR = 0.050); IL-1β, IL-6, and TNFα did not differ significantly from vehicle. At 24 h, no treatment differed significantly from vehicle in texture complexity. These results demonstrate that fine-grained texture features capture early stimulus-specific effects that are not fully resolved by coarse state assignment alone.

### 6. IFNγ specifically elevates intracellular ROS and correlates with the Spread-state shift

The morphological analyses in Fig. 4–5 established that each inflammatory stimulus drives a distinct pattern of iMGL remodeling, with IFNγ producing the largest redistribution toward the Spread state. To determine whether these morphological shifts are accompanied by altered functional output, we measured reactive oxygen species (ROS) production after 24 h inflammatory stimulation using CellROX deep-red staining (Fig. 6A). Morphological states were assigned to each cell in the ROS experiment using the frozen anchor coordinates defined in Fig. 3, ensuring a consistent single-cell state framework across experiments.

Among the six conditions, only IFNγ produced a significant increase in per-cell median CellROX intensity relative to vehicle (Fig. 6B; *p* < 0.01). LPS, IL-1β, IL-6, and TNFα did not significantly alter ROS levels at 24 h, despite driving measurable morphological changes (Fig. 4, 5). IFNγ was therefore both the strongest morphological perturbation and the only stimulus to elevate ROS at this time point.

Because IFNγ preferentially enriched the Spread state (Fig. 5), we next asked whether its effect on ROS was restricted to or concentrated in specific morphological states. Resolving per-cell ROS by state revealed that IFNγ-induced ROS elevation was broadly distributed: fold changes relative to vehicle were similar across all four states (all *p* > 0.05 after BH-FDR correction) (Fig. 6C). Thus, IFNγ elevates ROS cell-wide rather than through a state-specific subpopulation.

Finally, to test whether morphological remodeling and ROS production are coupled across stimuli, we correlated the per-experiment Spread-state fraction with median CellROX intensity (Fig. 6D). The two variables showed a positive trend across all treatment × experiment combinations (Spearman ρ = 0.42, p = 0.086, n = 18), with IFNγ occupying the upper-right region (highest Spread fraction and highest ROS). Although not statistically significant at the 0.05 level, the trend is consistent with the interpretation that the morphological and functional effects of IFNγ are coordinated, rather than independent consequences of inflammatory stimulation. Together, these results identify IFNγ as the only stimulus that combines the largest morphological shift with detectable ROS elevation among the six conditions tested, while indicating that ROS upregulation is not confined to a single morphological state.

## Discussion

Microglial morphology has long been used as a readout of activation or stimulus-dependent effects, but most studies still rely on relatively coarse classifications that do not fully capture the dynamic and stimulus-specific nature of the response. Here, we show that label-free, time-resolved single-cell profiling resolves microglial activation as graded shifts within a continuous morphological landscape, with each inflammatory stimulus occupying a distinct region of this landscape and following its own temporal trajectory. Moreover, we showed the morphological shift happened in parallel with a measurable change in redox state.

A key strength of this approach is the ability to resolve the temporal structure of microglial responses. Time-lapse profiling revealed that morphological divergence between control and co-stimulation with LPS and IFNγ emerges rapidly, peaks within hours, and subsequently attenuates over time. This transient trajectory, together with increased heterogeneity at intermediate time points, suggests that microglial activation involves dynamic and reversible remodeling rather than a stable endpoint state. This behavior is consistent with the increasingly recognized view that microglial activation is highly dynamic and context-dependent rather than confined to fixed activation states[29-31].

At the single-cell level, we identified four morphological states that capture the major axes of variation in iMGL shape. These states provide an interpretable framework for summarizing population-level responses and indicate that inflammatory stimulation primarily drives redistribution across pre-existing morphological configurations rather than the emergence of entirely new phenotypes. However, our results also show that discrete state assignments alone are insufficient to capture the full structure of the data. Multiple inflammatory stimuli induced broadly similar shifts in state composition, yet density-based analyses revealed distinct spatial distributions within the shared embedding. IFNγ, for example, produced a pronounced redistribution toward a specific subregion of the landscape, whereas IL-1β produced minimal changes at either the composition or density level. Together, these observations support a model in which microglial activation is better described as graded occupancy of a continuous morphological landscape than as transitions between mutually exclusive states[32-34].

In addition to spatial redistribution within the embedding, texture complexity provided an independent layer of information that further distinguished stimulus-specific responses. At early time points, LPS produced a significant reduction in texture complexity, with IFNγ showing a borderline reduction; TNFα and IL-1β produced smaller changes, and IL-6 was unchanged. Because these features capture sub-cellular organization that is invisible to shape-based descriptors, they demonstrate that stimulus identity is encoded simultaneously in coarse shape, spatial position within the morphological landscape, and fine-grained sub-cellular texture.

Linking morphology to function, we found that IFNγ, the cytokine producing the largest morphological redistribution, was also the only stimulus to elevate intracellular ROS at 24 h, and per-experiment Spread-state enrichment and median ROS showed a positive trend across treatments (Spearman *ρ* = 0.42, *p* = 0.086), with IFNγ producing both the largest morphological shift and the highest ROS levels. The IFNγ-induced ROS increase was broadly distributed across all four morphological states rather than concentrated in any single subpopulation, arguing against a model in which one discrete state accounts for the functional response. Instead, the coupling between morphological and functional remodeling appears to operate at the level of graded cellular states, reinforcing the value of continuous over categorical descriptors.

The hierarchy of responses across inflammatory stimuli broadly matches their established roles in microglial biology. IFNγ, which elicited the strongest and most distinct morphological redistribution, is a central mediator of classical inflammatory activation[35]. In contrast, IL-1β showed minimal effects in our analysis, suggesting that additional contextual signals may be required to drive robust morphological changes. Consistent with this, recent *in vivo* studies suggest that IL-1β acts primarily as a downstream amplifier of inflammatory cascades, indicating that its functional impact may originate more from autocrine/paracrine signaling within tissue contexts than from direct morphological responses to exogenous IL-1β[36]. The partial and overlapping responses observed for LPS, IL-6, and TNFα further suggest that microglial activation reflects integration of multiple signaling inputs into a shared phenotypic space.

More broadly, this work highlights the potential of label-free morphological profiling as a scalable and information-rich readout of cellular state, consistent with previous studies demonstrating that high-dimensional features extracted from cell images can serve as robust phenotypic fingerprints of biological perturbations[37]. Recent endpoint high-throughput, single-cell imaging-based approaches have applied multi-feature extraction and unsupervised clustering to characterize microglial morphological states, emphasizing the substantial heterogeneity of microglial phenotypes beyond simple binary classifications[38]. In contrast to these fluorescence-based and largely endpoint-driven approaches, our label-free framework enables continuous, non-perturbative monitoring of live cells. Therefore, this analysis allows for extending morphological profiling to capture dynamic and longitudinal changes in the cellular microglial state.

Future studies combining morphological profiling with molecular and functional readouts may enable a more comprehensive mapping of the microglial response landscape and its regulation in disease contexts. In addition, the scalability of this approach makes it well suited for therapeutic discovery, high-throughput perturbation and drug screening applications[39]. Together, our findings establish a framework for interpreting microglial morphology as a dynamic and continuous phenotype and provide a basis for quantitatively dissecting inflammatory responses at single-cell resolution.

## Limitations

Several limitations should be considered. First, morphological features provide an indirect readout of cellular state and do not directly capture the underlying molecular programs. Integrating morphological profiling with transcriptomic or proteomic measurements will be important to establish mechanistic links between morphology and function. Second, although steps were taken to control for technical variability, segmentation accuracy and imaging conditions may influence feature extraction, particularly for cells with complex or overlapping morphologies. Continued improvements in segmentation algorithms and quality control strategies will further enhance the robustness of this approach. Third, our analyses are based on three independent differentiation runs (n = 3). While experiment-level testing avoids pseudoreplication, this sample size limits statistical power, and several biologically meaningful effect-size shifts in state composition did not pass FDR correction (Table S2). Effect-size estimates (Δ fraction, Cohen’s D) and trends across stimuli therefore provide a more robust comparison than *p*-values alone, and the framework would benefit from larger biological-replicate cohorts in future applications. Finally, this study was performed using *in vitro* iPSC-derived microglia. While these models recapitulate key aspects of microglial biology, the extent to which the observed morphological landscapes reflect *in vivo* microglial states remains to be determined[40]. Extending this framework to more complex systems, including organoids[41] or *in vivo* imaging datasets, will be an important next step.

## Declarations

### Ethics approval and consent to participate

Not applicable. This study used human iPSC-derived cells *in vitro* and did not involve human participants, primary human tissue, or animal experiments.

### Consent for publication

Not applicable.

### Authors’ contributions

TC designed and performed experiments, developed the analysis pipeline, analyzed data, and wrote the manuscript. XL contributed to the initial conceptualization of the study and validated the analysis code. AMD supervised the study, acquired funding, and revised the manuscript. All authors read and approved the final manuscript.

## Acknowledgments

We also thank Prof. Bart Eggen (Department of Biomedical Sciences of Cells & Systems, University Medical Center Groningen) for generously providing the human iPSC line used in this study.

## Data availability

Raw phase-contrast images, Cellpose-SAM segmentation masks, CellProfiler per-cell measurement tables, plate-level metadata files, and other data required to reproduce the analyses are available on Zenodo at https://doi.org/10.5281/zenodo.19826203. The repository README and data/README.md describe the expected local directory structure for reproducing the analysis.

## Use of artificial intelligence

AI-based tools were used to assist with code and text refinement. All analyses were performed and validated by the authors.

---

**Figure S1.**
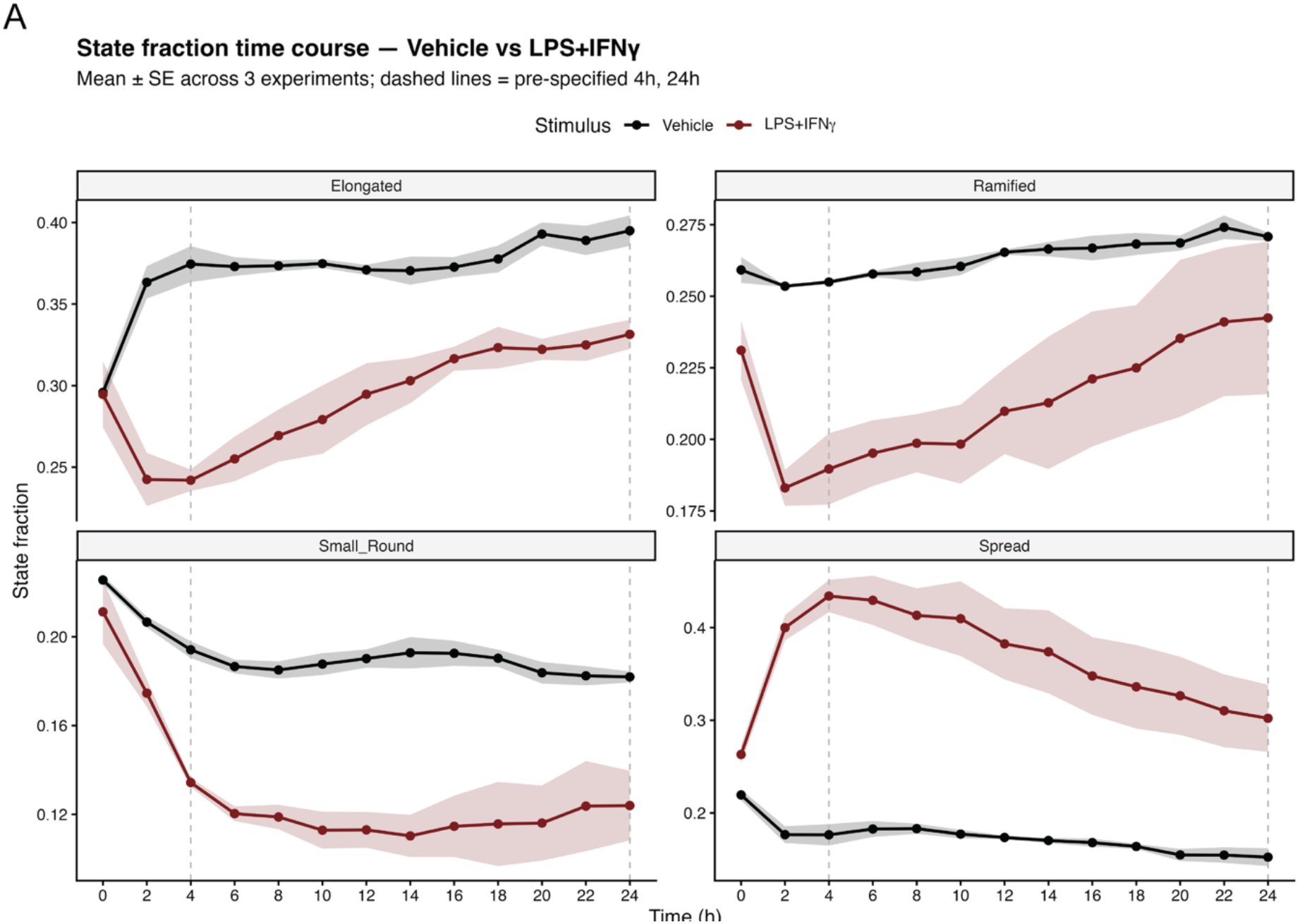
State fraction time course — vehicle vs LPS+IFNγ. Fraction of iMGLs in each of the four morphological states across 13 time points over 24 h. Each cell was assigned to its nearest anchor using the frozen classifier from Fig. 3. Lines show mean ± SE across n = 3 independent experiments; dashed lines mark the pre-specified time points (4 h, 24 h) used in Fig. 3.

**Figure S2.**
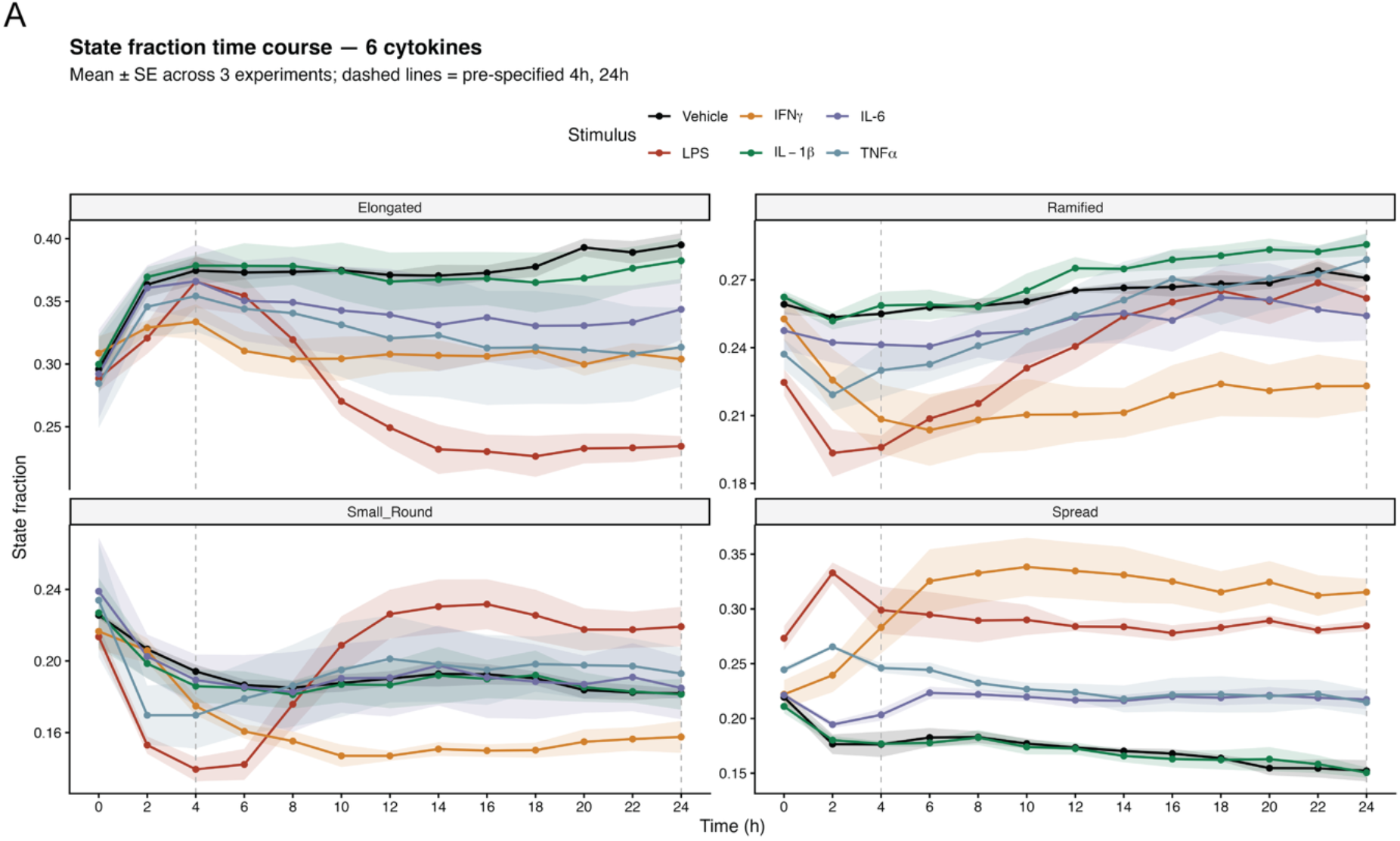
State fraction time course — five inflammatory stimuli. Fraction of iMGLs in each of the four morphological states across 13 time points over 24 h, shown for vehicle, LPS, IFNγ, IL-1β, IL-6, and TNFα. Each cell was assigned to its nearest anchor using the frozen classifier from Fig. 3. Lines show mean ± SE across n = 3 independent experiments; dashed lines mark the pre-specified time points (4 h, 24 h) used in Fig. 5.

**Figure S3.**
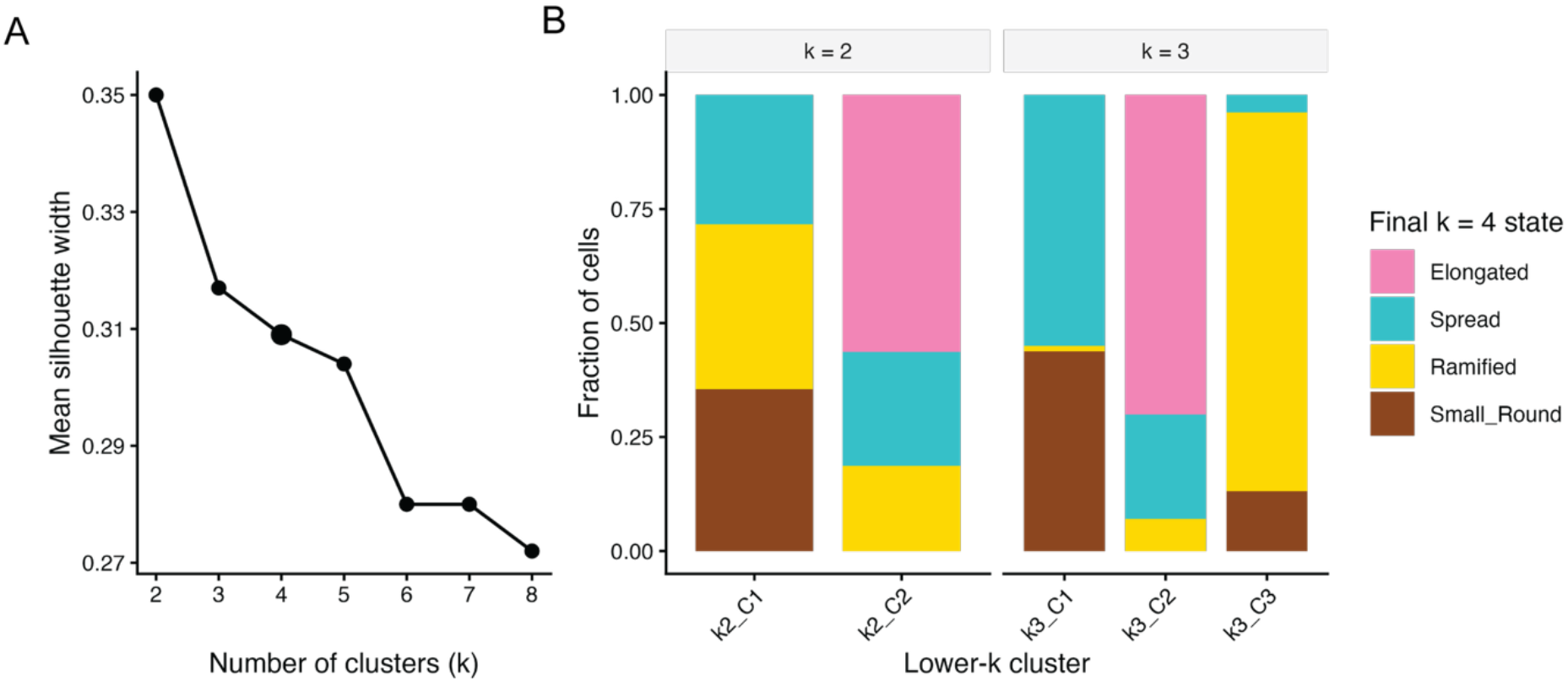
Sensitivity analysis for the number of morphological anchor states. (A) Mean silhouette width for k = 2–8 computed on a 5,000-cell random subsample of the fixed training set in the three-axis anchor space (z_size, z_round, z_elong). The final k = 4 solution is highlighted. (B) Cross-tabulation of the final k = 4 anchor states against k = 2 and k = 3 clustering solutions on the same subsample. Lower-k solutions merged multiple final anchor states, whereas k = 4 provided interpretable and reusable anchor states corresponding to Elongated, Spread, Ramified, and Small_Round morphologies.

## References

[1] F. Ginhoux, M. Greter, M. Leboeuf, S. Nandi, P. See, S. Gokhan, M.F. Mehler, S.J. Conway, L.G. Ng, E.R. Stanley, I.M. Samokhvalov, M. Merad, Fate mapping analysis reveals that adult microglia derive from primitive macrophages, Science 330(6005) (2010) 841–5.

[2] L.C. Mehl, A.V. Manjally, O. Bouadi, E.M. Gibson, T.L. Tay, Microglia in brain development and regeneration, Development 149(8) (2022).

[3] C.N. Parkhurst, G. Yang, I. Ninan, J.N. Savas, J.R. Yates, 3rd, J.J. Lafaille, B.L. Hempstead, D.R. Littman, W.B. Gan, Microglia promote learning-dependent synapse formation through brain-derived neurotrophic factor, Cell 155(7) (2013) 1596–609.

[4] Y. Hattori, The multifaceted roles of embryonic microglia in the developing brain, Front Cell Neurosci 17 (2023) 988952.

[5] A. Ajoolabady, B. Kim, A.A. Abdulkhaliq, J. Ren, S. Bahijri, J. Tuomilehto, A. Borai, J. Khan, D. Pratico, Dual role of microglia in neuroinflammation and neurodegenerative diseases, Neurobiol Dis 216 (2025) 107133.

[6] W. Gomes-Leal, Microglial physiopathology: how to explain the dual role of microglia after acute neural disorders?, Brain Behav 2(3) (2012) 345–56.

[7] E.A. Ling, W.C. Wong, The origin and nature of ramified and amoeboid microglia: a historical review and current concepts, Glia 7(1) (1993) 9–18.

[8] L.J. Lawson, V.H. Perry, P. Dri, S. Gordon, Heterogeneity in the distribution and morphology of microglia in the normal adult mouse brain, Neuroscience 39(1) (1990) 151–70.

[9] A. Vidal-Itriago, R.A.W. Radford, J.A. Aramideh, C. Maurel, N.M. Scherer, E.K. Don, A. Lee, R.S. Chung, M.B. Graeber, M. Morsch, Microglia morphophysiological diversity and its implications for the CNS, Front Immunol 13 (2022) 997786.

[10] M. Augusto-Oliveira, G.P. Arrifano, C.G. Leal-Nazaré, A. Chaves-Filho, L. Santos-Sacramento, A. Lopes-Araujo, M. Tremblay, M.E. Crespo-Lopez, Morphological diversity of microglia: Implications for learning, environmental adaptation, ageing, sex differences and neuropathology, Neurosci Biobehav Rev 172 (2025) 106091.

[11] E.M. Abud, R.N. Ramirez, E.S. Martinez, L.M. Healy, C.H.H. Nguyen, S.A. Newman, A.V. Yeromin, V.M. Scarfone, S.E. Marsh, C. Fimbres, C.A. Caraway, G.M. Fote, A.M. Madany, A. Agrawal, R. Kayed, K.H. Gylys, M.D. Cahalan, B.J. Cummings, J.P. Antel, A. Mortazavi, M.J. Carson, W.W. Poon, M. Blurton-Jones, iPSC-Derived Human Microglia-like Cells to Study Neurological Diseases, Neuron 94(2) (2017) 278-293.e9.

[12] E. Wogram, F. Sümpelmann, A. Khalil, A. Flamier, D. Fu, G.W. Bell, R. Jaenisch, Human iPSC-Derived Microglia Integrate Into Cerebral Organoids and Assume an In Vivo-Like Phenotype, Eur J Neurosci 62(9) (2025) e70281.

[13] P. Douvaras, B. Sun, M. Wang, I. Kruglikov, G. Lallos, M. Zimmer, C. Terrenoire, B. Zhang, S. Gandy, E. Schadt, D.O. Freytes, S. Noggle, V. Fossati, Directed Differentiation of Human Pluripotent Stem Cells to Microglia, Stem Cell Reports 8(6) (2017) 1516–1524.

[14] O. Uriarte Huarte, L. Richart, M. Mittelbronn, A. Michelucci, Microglia in Health and Disease: The Strength to Be Diverse and Reactive, Front Cell Neurosci 15 (2021) 660523.

[15] J. Leyh, S. Paeschke, B. Mages, D. Michalski, M. Nowicki, I. Bechmann, K. Winter, Classification of Microglial Morphological Phenotypes Using Machine Learning, Front Cell Neurosci 15 (2021) 701673.

[16] Y. Ding, M.C. Pardon, A. Agostini, H. Faas, J. Duan, W.O.C. Ward, F. Easton, D. Auer, L. Bai, Novel Methods for Microglia Segmentation, Feature Extraction, and Classification, IEEE/ACM Trans Comput Biol Bioinform 14(6) (2017) 1366–1377.

[17] K. Chen, X. Qi, L.L. Zhu, M.L. Li, B. Cong, Y.M. Li, Quantitative analysis of microglia morphological changes in the hypothalamus of chronically stressed rats, Brain Res Bull 206 (2024) 110861.

[18] B. Cătălin, L. Stopper, T.A. Bălşeanu, A. Scheller, The in situ morphology of microglia is highly sensitive to the mode of tissue fixation, J Chem Neuroanat 86 (2017) 59–66.

[19] K.E. Hopperton, D. Mohammad, M.O. Trépanier, V. Giuliano, R.P. Bazinet, Markers of microglia in post-mortem brain samples from patients with Alzheimer’s disease: a systematic review, Mol Psychiatry 23(2) (2018) 177–198.

[20] K. Kierdorf, D. Erny, T. Goldmann, V. Sander, C. Schulz, E.G. Perdiguero, P. Wieghofer, A. Heinrich, P. Riemke, C. Hölscher, D.N. Müller, B. Luckow, T. Brocker, K. Debowski, G. Fritz, G. Opdenakker, A. Diefenbach, K. Biber, M. Heikenwalder, F. Geissmann, F. Rosenbauer, M. Prinz, Microglia emerge from erythromyeloid precursors via Pu.1- and Irf8-dependent pathways, Nat Neurosci 16(3) (2013) 273–80.

[21] R.L. Lowery, M.E. Tremblay, B.E. Hopkins, A.K. Majewska, The microglial fractalkine receptor is not required for activity-dependent plasticity in the mouse visual system, Glia 65(11) (2017) 1744–1761.

[22] E.A. Specht, E. Braselmann, A.E. Palmer, A Critical and Comparative Review of Fluorescent Tools for Live-Cell Imaging, Annu Rev Physiol 79 (2017) 93–117.

[23] E.C. Jensen, Use of fluorescent probes: their effect on cell biology and limitations, Anat Rec (Hoboken) 295(12) (2012) 2031–6.

[24] M. Pachitariu, M. Rariden, C. Stringer, Cellpose-SAM: superhuman generalization for cellular segmentation, bioRxiv (2025) 2025.04.28.651001.

[25] D.R. Stirling, M.J. Swain-Bowden, A.M. Lucas, A.E. Carpenter, B.A. Cimini, A. Goodman, CellProfiler 4: improvements in speed, utility and usability, BMC Bioinformatics 22(1) (2021) 433.

[26] N. Fattorelli, A. Martinez-Muriana, L. Wolfs, I. Geric, B. De Strooper, R. Mancuso, Stem-cell-derived human microglia transplanted into mouse brain to study human disease, Nat Protoc 16(2) (2021) 1013–1033.

[27] A.M. Sabogal-Guáqueta, A. Marmolejo-Garza, M. Trombetta-Lima, A. Oun, J. Hunneman, T. Chen, J. Koistinaho, S. Lehtonen, A. Kortholt, J.C. Wolters, B.M. Bakker, B.J.L. Eggen, E. Boddeke, A. Dolga, Species-specific metabolic reprogramming in human and mouse microglia during inflammatory pathway induction, Nature Communications 14(1) (2023) 6454.

[28] M. Koskuvi, E. Pörsti, T. Hewitt, N. Räsänen, Y.C. Wu, K. Trontti, A. McQuade, S. Kalyanaraman, I. Ojansuu, O. Vaurio, T.D. Cannon, J. Lönnqvist, S. Therman, J. Suvisaari, J. Kaprio, M. Blurton-Jones, I. Hovatta, M. Lähteenvuo, T. Rolova, Š. Lehtonen, J. Tiihonen, J. Koistinaho, Genetic contribution to microglial activation in schizophrenia, Mol Psychiatry 29(9) (2024) 2622–2633.

[29] N. Javanmehr, K. Saleki, P. Alijanizadeh, N. Rezaei, Microglia dynamics in aging-related neurobehavioral and neuroinflammatory diseases, J Neuroinflammation 19(1) (2022) 273.

[30] N. Stence, M. Waite, M.E. Dailey, Dynamics of microglial activation: a confocal time-lapse analysis in hippocampal slices, Glia 33(3) (2001) 256–66.

[31] B.M. Davis, M. Salinas-Navarro, M.F. Cordeiro, L. Moons, L. De Groef, Characterizing microglia activation: a spatial statistics approach to maximize information extraction, Sci Rep 7(1) (2017) 1576.

[32] R. Martins-Ferreira, J. Calafell-Segura, B. Leal, J. Rodríguez-Ubreva, E. Martínez-Saez, E. Mereu, E.C.P. Pinho, A. Laguna, E. Ballestar, The Human Microglia Atlas (HuMicA) unravels changes in disease-associated microglia subsets across neurodegenerative conditions, Nat Commun 16(1) (2025) 739.

[33] M. Olah, V. Menon, N. Habib, M.F. Taga, Y. Ma, C.J. Yung, M. Cimpean, A. Khairallah, G. Coronas-Samano, R. Sankowski, D. Grün, A.A. Kroshilina, D. Dionne, R.A. Sarkis, G.R. Cosgrove, J. Helgager, J.A. Golden, P.B. Pennell, M. Prinz, J.P.G. Vonsattel, A.F. Teich, J.A. Schneider, D.A. Bennett, A. Regev, W. Elyaman, E.M. Bradshaw, P.L. De Jager, Single cell RNA sequencing of human microglia uncovers a subset associated with Alzheimer’s disease, Nat Commun 11(1) (2020) 6129.

[34] T.R. Hammond, C. Dufort, L. Dissing-Olesen, S. Giera, A. Young, A. Wysoker, A.J. Walker, F. Gergits, M. Segel, J. Nemesh, S.E. Marsh, A. Saunders, E. Macosko, F. Ginhoux, J. Chen, R.J.M. Franklin, X. Piao, S.A. McCarroll, B. Stevens, Single-Cell RNA Sequencing of Microglia throughout the Mouse Lifespan and in the Injured Brain Reveals Complex Cell-State Changes, Immunity 50(1) (2019) 253-271.e6.

[35] O. Kann, F. Almouhanna, B. Chausse, Interferon γ: a master cytokine in microglia-mediated neural network dysfunction and neurodegeneration, Trends Neurosci 45(12) (2022) 913–927.

[36] A. Grayston, M. Baptista, K. Wemyss, R. Taylor, G. Cullen, S.S. Jafree, N. Luka, J.R. Cox, J.E. Konkel, D. Brough, S.M. Allan, E. Pinteaux, Specific deletion of interleukin-1 beta in microglia improves acute outcome and modulates neurogenesis after ischemic stroke, bioRxiv (2025) 2025.12.19.695391.

[37] M.A. Bray, S. Singh, H. Han, C.T. Davis, B. Borgeson, C. Hartland, M. Kost-Alimova, S.M. Gustafsdottir, C.C. Gibson, A.E. Carpenter, Cell Painting, a high-content image-based assay for morphological profiling using multiplexed fluorescent dyes, Nat Protoc 11(9) (2016) 1757–74.

[38] J. Kim, P. Pavlidis, A.V. Ciernia, Development of a High-Throughput Pipeline to Characterize Microglia Morphological States at single-cell resolution, eNeuro 11(7) (2024).

[39] A.M. Dolga, C. Culmsee, Protective Roles for Potassium SK/K(Ca)2 Channels in Microglia and Neurons, Front Pharmacol 3 (2012) 196.

[40] A.M. Sabogal-Guáqueta, A. Marmolejo-Garza, V.P. de Pádua, B. Eggen, E. Boddeke, A.M. Dolga, Microglia alterations in neurodegenerative diseases and their modeling with human induced pluripotent stem cell and other platforms, Prog Neurobiol 190 (2020) 101805.

[41] A.M. Sabogal-Guaqueta, T. Mitchell-Garcia, J. Hunneman, D. Voshart, A. Thiruvalluvan, F. Foijer, F. Kruyt, M. Trombetta-Lima, B.J.L. Eggen, E. Boddeke, L. Barazzuol, A.M. Dolga, Brain organoid models for studying the function of iPSC-derived microglia in neurodegeneration and brain tumours, Neurobiol Dis 203 (2024) 106742.

